# Epstein-Barr Virus May Contribute to Central Nervous System Involvement in HIV-positive Individuals

**DOI:** 10.1101/341354

**Authors:** T Lupia, MG Milia, C Atzori, S Audagnotto, D Imperiale, L Mighetto, V Pirriatore, G Gregori, F Lipani, V Ghisetti, S Bonora, G Di Perri, A Calcagno

## Abstract

Epstein-Barr virus (EBV) often accesses the central nervous system (CNS) where it may lead to blood brain barrier (BBB) integrity disruption, facilitating the migration of immune cells into brain parenchyma. Our aim was to study the association between cerebrospinal fluid (CSF) EBV DNA and HIV-1 compartmental replication. 281 HIV-positive adults undergoing lumbar punctures for clinical reasons (excluding those with lymphoproliferative disorders) and CSF samples were examined. CSF virological, neurodamage (tau, p-tau, 1-42 beta amyloid) and immune activation (neopterin and S100beta) markers were measured by immune-enzymatic, ELISA and PCR validated methods. Two hundred eighty one patients were included; 111 (40.5 %) were naïve for antiretroviral treatment. CSF EBV DNA was detectable in 25 (21.9%) naïve and 26 (16%) treated patients at low levels (<100 and 146 copies/mL). Naïve EBV+ subjects presented higher CSF HIV RNA, biomarkers (t-tau, p-tau, neopterin) and higher rates of pleocytosis. Treated EBV+ individuals showed pleocytosis, higher CSF HIV RNA, CSF to serum albumin ratio, IgG index and neopterin. No association was observed between detectable CSF EBV DNA and the rate of CSF escape. In patients with plasma HIV RNA <20 copies/mL (n=97) CSF EBV DNA was detectable in 13 subjects (13.4%) and it was associated with pleocytosis, higher CSF HIV RNA and neopterin levels. EBV DNA was detectable in a considerable proportion of HIV-positive patients and it was associated with higher levels of CSF HIV RNA and neuronal damage/inflammation biomarkers. The role of EBV reactivation in HIV-associated CNS disorders warrant further studies.

**Importance:** EBV is a human gamma-herpesvirus with a seroprevalence in adults approaches 95% and the pattern of clinical manifestations is very heterogeneous and varies from asymptomatic or mild viral infection to a tightly linked with several malignancies as nasopharyngeal carcinoma, Hodgkin’s lymphoma and Burkitt’s lymphoma. HIV-infected and immunocompetent patients were both at risk of primary infection and complications linked to EBV.

Primary tropism of EBV is for lymphocytes (type B, T and NK), epithelial, endothelial and smooth muscle cells and establishes lifelong latent infection. Central nervous system could be affected by this herpesvirus in primary infection and reactivation and EBV-DNA is not an uncommon finding in CSF in HIV-infected population. The significance of our research is in identifying the presence of a link between HIV and EBV CNS replication.

## Introduction

Epstein-Barr virus (EBV) or Human Herpesvirus 4 (HHV-4) is a widely disseminated gamma herpesvirus capable to persist, lifelong and asymptomatically, in a latent infection in adults [1]. Transmission of EBV is so efficient that by adulthood most (> 95%) of the world’s population has been infected [2].

EBV may affect the central nervous system (CNS) and clinical manifestations were first noted in 1931 by Epstein and Johansen. Both primary infection and reactivation can cause neurological diseases and central nervous system involvement occurs in 1 to 10% of the cases [3]. Individual state of immunocompetence, age and comorbidities have been associated with the occurrence of neurological complications that include: meningo-encephalites, cerebellitis, optic neuritis, cranial nerve palsy, peripheral neuropathy, Alice in Wonderland syndrome, ataxia, chorea, post-infectious autoimmune disorders, including Guillain-Barrè syndrome, acute disseminated encephalomyelitis (ADEM) and transverse myelitis [3][4][5].

Biopsy proven vasculitis due to EBV infection have been reported: perivascular inflammatory infiltrate was dominantly composed of CD3+ and CD8+ T-lymphocytes and macrophages. Some of the CD3 positive cells were also EBV-encoded RNA-1 (EBER1) positive, one of the two small noncoding RNAs (EBER1 & 2) found in latently EBV infected cells [6][7][8].

In MS patients’ brains, EBV infection in B cells seems to alters the ability of B cells to process and present a pathogenetically relevant myelin autoantigen and expression of higher levels of costimulatory molecules than healthy controls, suggesting an enhanced APC function of B cells in MS brains, leading to an higher autoimmune risk [9]. EBV-positive B lymphocytes count in normal human brain is very low, but is shown an higher cell number in HIV infected brains. In PCR-based studies and in situ hybridation studies were shown a detection of EBV in both lymphomatoid tissue and in pleomorphic lymphoid infiltrates [10].

Several works have established that EBV can infect macrovascular endothelial cells in human tissue [11] [12], human brain microvessels [13] and in culture with human umbilical vein endothelial cells (HUVECs) [14] [15]. In endothelial cells with lytic reactivation of EBV were found an increase production of pro- inflammatory molecules (CCL-2 and CCL-5) and also hyper-expression of adhesion molecules on surfaces (ICAM-1 and VCAM-1) with a potential creation of an inflammatory breach through the Blood Brain Barrier (BBB) [13] [14] [15] [16]

HIV-positive patients have a higher risk of EBV-associated diseases due to reduced immune surveillance; non-Hodgkin and Burkitt’s lymphomas are among AIDS-defining conditions. Highly Active Antiretroviral Treatment (HAART) has dramatically reduced the incidence of HIV-associated dementia but milder forms of cognitive impairment as well as cerebrospinal fluid (CSF) HIV escaper persist despite treatment. Beside incomplete CSF antiretrovirals’ penetration/effectiveness HIV escape may be due to enhanced blood brain barrier (BBB) permeability and secondary to other concomitant infections (“secondary escape”) [17]. We aimed at studying the role of CSF EBV DNA in HIV-positive subjects in terms of HIV replication, BBB damage and biomarkers of neuronal damage/inflammation.

## Material and Methods

Adult HIV-positive patients undergoing lumbar punctures for clinical reasons, were enrolled. Patients with primary central nervous system lymphomas (PCNSLs), lymphoproliferative diseases (Lds) and autoimmune disorders were excluded. Demographic, immunovirological, clinical and therapeutic data were recorded as well as CSF features. The protocol was approved by our Ethics Committee (Comitato Etico Interaziandale di Orbassano, n. 103/2015). Patients signed a written informed consent at enrollment.

HIV RNA was measured through the real time Polymerase Chain Reaction (PCR) assay CAP/CTM HIV-1 vs. 2.0 (CAP/CTM, Roche Molecular System, Branchburg,NJ, detection limit: 20 copies/mL of HIV-1 RNA). EBV DNA was measured through the real time Polymerase Chain Reaction (PCR) (detection limit: 100 copies/mL of EBV DNA).

Quantitative determination of albumin in serum and CSF was measured by Immunoturbidimetric methods (AU 5800, Beckman Coulter, Brea, CA, USA). CSAR, calculated as CSF albumin (mg/L)/serum albumin (g/L), was used to evaluate BBB function. BBB damage definition was derived from age-adjusted Reibergrams (normal if below 6.5 in patients aged <40 years and below 8 in patients >40 years).

CSF total tau (t-tau), phosphorylated tau (p-tau) and β-amyloid1-42 (Aβ1-42) were measured by immunoenzymatic methods (Fujirebio diagnostics, Malvern, U.S.A.) with limits of detection respectively of 57, 20 and 225 pg/ml. Neopterin and S100B were measured through validated ELISA methods [DRG Diagnostics (Marburg, Germany) and DIAMETRA S.r.l. (Spello, Italy), respectively]. Reference values were as follows: t-tau [<300 pg/mL (in patients aged 21–50), <450 pg/mL (in patients aged 51–70) or <500 pg/mL in older patients], p-tau (<61 pg/mL), 1–42 beta amyloid (>500 pg/mL), neopterin (<1.5 ng/mL) and S100B (<380 pg/mL) [18].

HIV RNA was measured through the real time Polymerase Chain Reaction (PCR) assay CAP/CTM HIV-1 vs. 2.0 (CAP/CTM, Roche Molecular System, Branchburg, NJ, detection limit: 20 copies/mL of HIV-1 RNA). EBV DNA was measured through the real time Polymerase Chain Reaction (PCR) (detection limit: 150 copies/mL of EBV DNA).

Data were analyzed using standard statistical methods: variables were described with medians [interquartile ranges (IQR) or ranges (minimum-maximum)] and they were compared using non-parametric tests (Mann–Whitney, Kruskal-Wallis and Spearman’s tests as specified in the text). Data analysis was performed using PASW software version 22.0 (IBM).

## Results

Two hundred and eighty one adult patients were included. 111 (40.5 %) patients were naïve for combination antiretroviral treatment (cART); baseline and immune-virological characteristics, stratified by cART use, are shown in Table 1. Lumbar punctures were performed before starting antiretroviral treatment in naïve late presenting subjects (CD4+ T lymphocytes <100/uL) or in symptomatic treated patients with cognitive disorders, headache or other neurological complaints.

**Table 1.** Baseline characteristics. Demographic, immunovirological and clinical variables of subjects at baseline. “BMI” body mass index, “VL” viral load, “CSF” cerebrospinal fluid, “HAART” Highly Active Antiretroviral Treatment, “NNRTI” non-nucleoside reverse transcriptase inhibitor, “PI” protease inhibitor, “INSTI” integrase strand-transfer inhibitor, “MVC” Maraviroc, “HAND” HIV-1 associated neurocognitive disorder.

|  | **Naive** | **Treated** |
| --- | --- | --- |
| **n** | 111 | 170 |
| **Age:** years | 44 (37.5-48) | 48 (42.3-56.1) |
| **Male gender:** n (%) | 78 (70.7%) | 117 (69.1%) |
| **BMI:** Kg/m<sup>2</sup> | 24.5 (19,6 – 25,1) | 22.8 (20-25.2) |
| **CD4:** Cells/uL | 67 (27 – 157) | 319 (116 – 570) |
| **CD4 nadir:** Cells/uL | 64 (25 – 141) | 46 (13 – 194) |
| **plasma VL:** Log<sub>10</sub> copies/mL | 5.3 (4.9 - 5.9) | 1.3 (<1.3 – 1.9) |
| **CSF VL:** Log<sub>10</sub> copies/mL | 3.9 (2.9 - 4.7) | 1.4 (<1.3 – 2.1) |
| **Time with HIV-positive test:** months | 78 (47 – 205) | 197 (95 – 290) |
| **ARV classes:** NNRTI | - | 43 |
| PI | - | 106 |
| INSTI | - | 45 |
| MVC | - | 16 |
| **Indication for LP:**Late Presentation | 48 | 49 |
| Opportunistic Infection | 24 | 32 |
| HAND | 16 | 40 |
| Syphilis | 1 | 5 |
| Follow up | 0 | 0 |
| Headache | 0 | 2 |
| **CSF EBV detectable:** n (%) | 25 (21.9%) |  |
| **CSF EBV DNA in those with detectable levels: copies/mL** | <100 (< 100 – 292) | 26 (16%)<br>141 (< 100 - 448) |

CSF EBV DNA was detectable in 25 naïve (21.9%) and 26 treated (16%) patients with median values of <100 (<100-234) and 146 (<100-612) copies/mL respectively. Virological, neuronal damage and inflammation biomarkers stratified by cART use and CSF EBV detection are shown in Table 2.

**Table 2.** Cerebrospinal fluid biomarkers characteristics of the four groups. Immunoactivation and neurodamage markers, stratified according to presence or not of EBV DNA on CSF and cART. “CSF” cerebrospinal fluid, “CSAR” Cerebrospinal fluid Serum Albumin Ratio, “BBBi” Blood Brain Barrier impairment, “t-tau” total tau, “p-tau”phosphorylated tau, “Aβ1-42” 1-42 beta amyloid, “S100B” S100 Beta. “n.s.” non significant (p values >0.05).

|  | Naive |  |  | Treated |  |  |
| --- | --- | --- | --- | --- | --- | --- |
|  | CSF EBV- | CSF EBV+ | P value | CSF EBV- | CSF EBV+ | P value |
| **CSF cells: n/uL** | 0 (0-3) | 10 (0-43) | <0.001 | 0 (0-3) | 0 (0-40) | <0.001 |
| **CSF pleocytosis** | 12.6% | 52% | <0.001 | 8.6% | 40% | <0.001 |
| **CSAR** | 5.9<br>(4 – 8.7) | 7.2<br>(4.1 – 11.2) | n.s. | 5.1<br>(3.7-7.0) | 6.1<br>(5.2-10.6) | 0.008 |
| **BBBi** | 41.2% | 47.8% | n.s. | 28.3% | 45.8% | 0.090 |
| **IgG index** | 0.4<br>(0.2-0.7) | 0.5<br>(0.2-0.8) | n.s. | 0.4<br>(0.2-0.6) | 0.5<br>(0.3-0.8) | 0.020 |
| **t-tau: pg/mL** | 114<br>(67.7-202) | 233<br>(130-334) | 0.001 | 130.1<br>(58-212) | 62<br>(37.5-153) | n.s. |
| **p-tau: pg/mL** | 29<br>(21.6-39,2) | 38<br>(31-44.7) | 0.034 | 35<br>(25-41.8) | 28.9<br>(21.5-37.5) | n.s. |
| **Aβ1-42: ng/mL** | 904<br>(643 – 1105) | 839<br>(584 – 1181) | n.s. | 893<br>(748-1075) | 788<br>(636-882) | 0.037 |
| **Neopterin: ng/mL** | 2.02 | 9.25 | <0.001 | 0.81 | 1.79 | 0.070 |
|  | (1.1 – 3.8) | (3.3-12.1) |  | (0.5-1.4) | (0.6-4.3) |  |
| **S100B:** pg/mL | 150.7<br>(87 - 242) | 188.4<br>(96 – 279) | n.s. | 140<br>(85-206) | 54<br>(37.5-210) | n.s. |

Naïve patients with detectable EBV DNA had higher CSF HIV viral load (4.2 vs. 3.7 log_10_ copies/mL, p=0.010); CSF to plasma HIV RNA ratios (25 vs. 4%, p=0.025), higher rates of pleocytosis (52% vs. 12.6%, p<0.001), CSF neuronal damage biomarkers t-tau (233 vs. 114 pg/mL, p=0.002), p-tau (38 vs. 29 pg/mL, p=0.051) and neopterin (9.25 vs. 2.02 ng/mL, p=0.001) (Figure 1 and 2, above).

**Figure 1:**
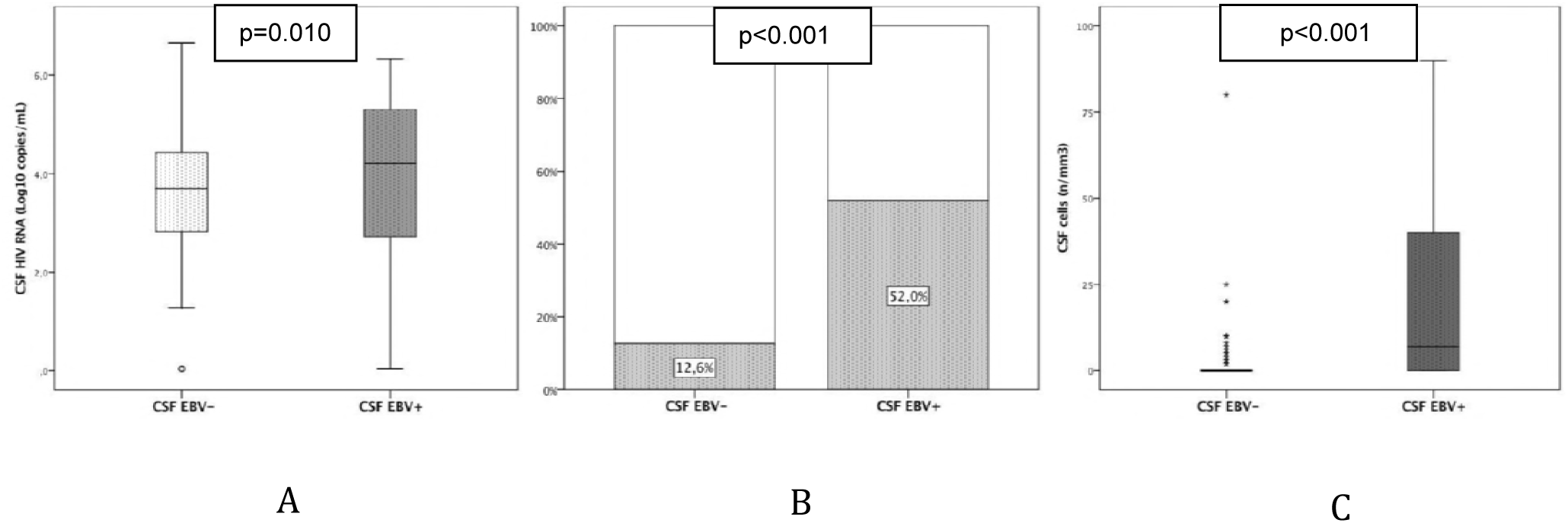
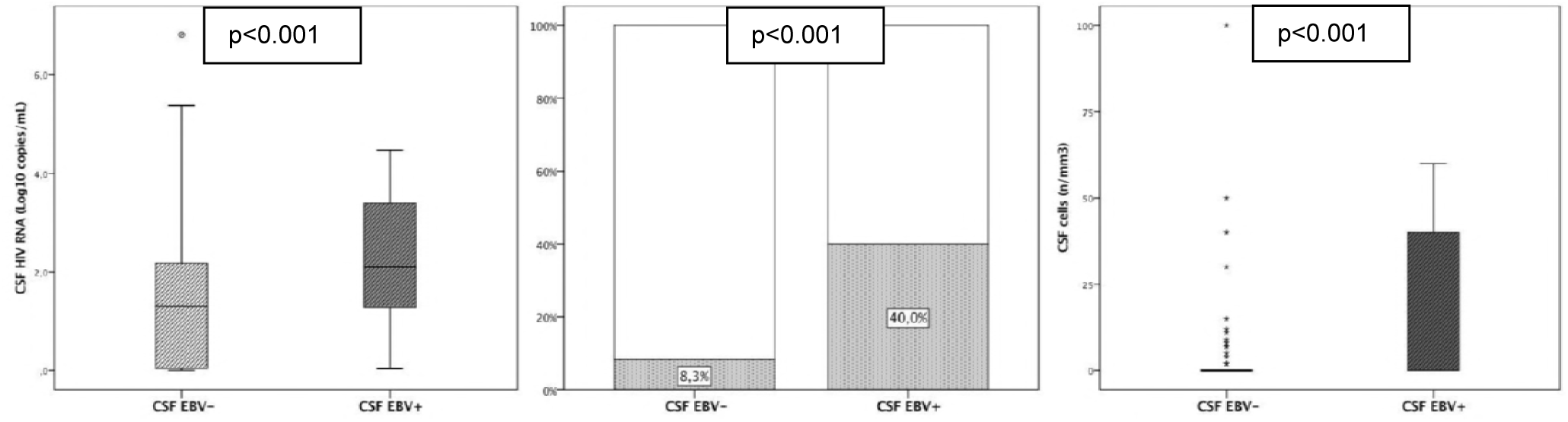
Cerebrospinal fluid HIV RNA (A), pleocytosis (B) and cell numbers (C) in naïve (above) and treated patients (below). “CSF”, cerebrospinal fluid; “CSF EBV+”, detectable CSF EBV DNA.

**Figure 2:**
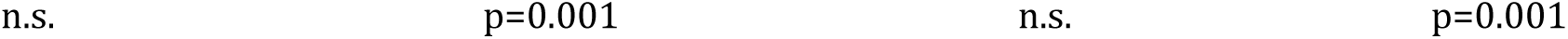
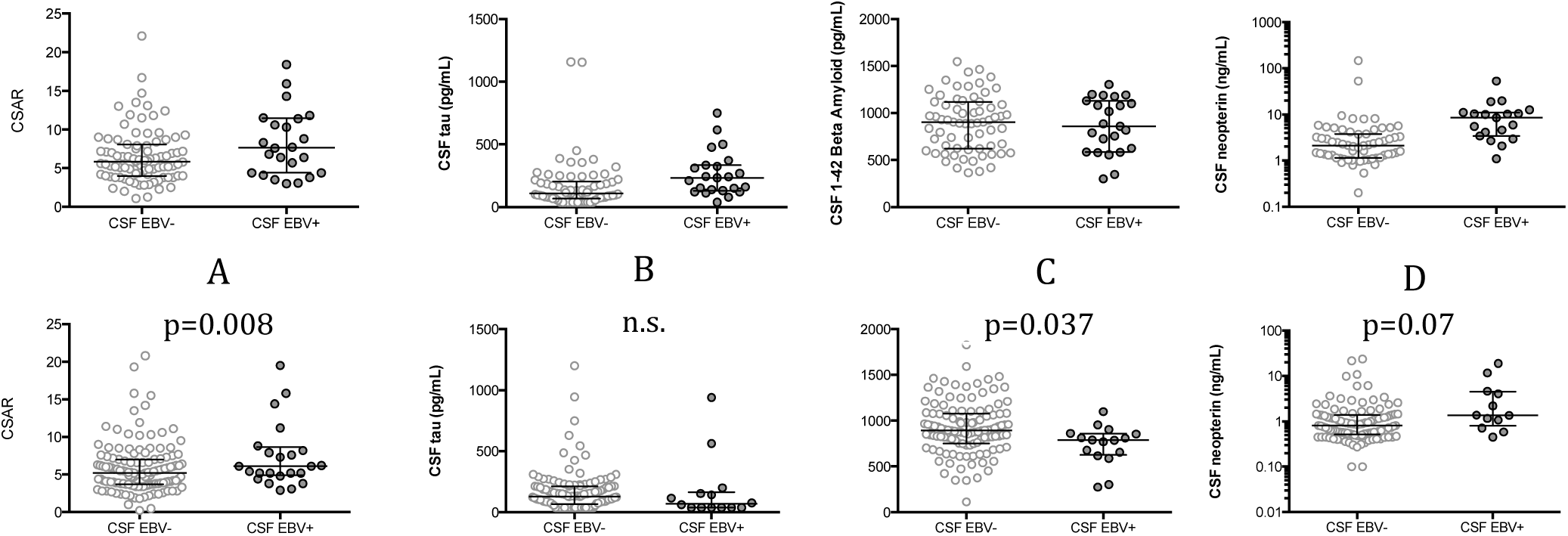
Cerebrospinal fluid biomarkers concentrations in naïve (above) and treated patients (below); cerebrospinal fluid to serum albumin ratio (A), total tau (B), 1-42 beta amyloid (C) and neopterin (D). “CSF”, cerebrospinal fluid; “CSF EBV+”, detectable CSF EBV DNA, “CSAR”; CSF to serum albumin ratio. All scatter dot plots present a central bar (median) with lateral error bars (IQR). In all graphs white dots represent observations in the EBV-negative group (left) instead black dots picture observations in EBV-positive group (right).

Treated patients with detectable EBV DNA had lower CD4 cell counts (136 vs. 287 cell/uL, p=0.015) and higher CSF HIV RNA (2.1 vs. <1.3 log_10_ copies/mL, p<0.001), higher levels of pleocytosis (40 vs. 8.3%, p<0.001), CSAR (6.1 vs. 5.1, p=0.011), IgG index (4.7 vs. 3.8, p=0.042) and neopterin (1.79 vs. 0.81 ng/mL, p=0.009) (Figure 1 and 2, below). Conversely 1-42 beta amyloid was lower in EBV-positive individuals (788 vs. 893 pg/mL, p=0.007).

The rate of CSF escape was similar in EBV-positive naïve (8.3% vs. 4.5%, p=0.607) and treated patients (28% vs. 23.3%, p=0.615).

In plasma controllers (HIV RNA <20 copies/mL, n=97) CSF EBV DNA was detectable in 13 individuals (13.4%): it was associated with pleocytosis (50% vs. 8.4%, p=0.001), higher CSF HIV RNA in those with detectable viral load (2.28 vs. 1.85 Log_10_ copies/mL, p=0.011) and higher CSF neopterin levels [2.81 ng/mL vs. 0.77 ng/mL, p=0.012] (Figure 3). In CSF controllers (CSF HIV RNA <20 copies/mL, n=84) CSF EBV DNA was detectable in 10 individuals (11.9%): it was associated with pleocytosis (66.6% vs. 6.9%, p<0.001) and border-line higher CSF neopterin (3.74 vs. 0.78, p=0.06) (Figure 3).

**Figure 3:**
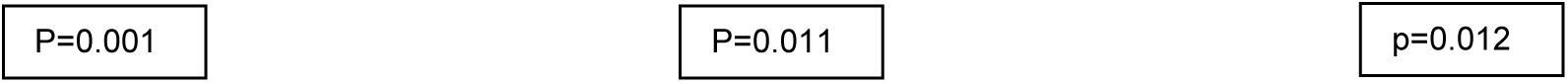
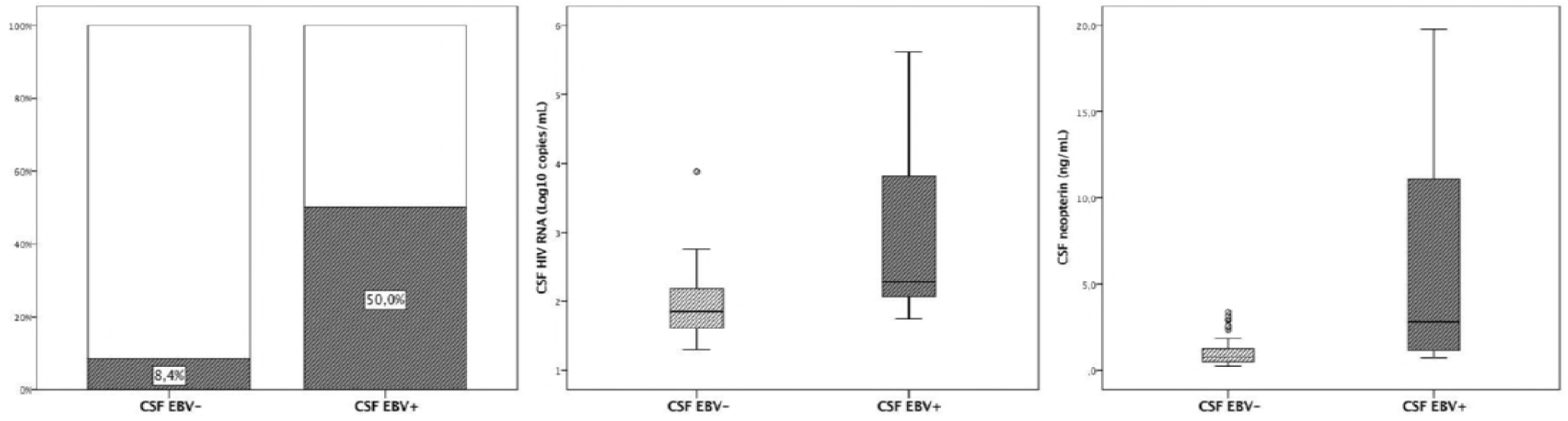
Cerebrospinal fluid pleocytosis (A), HIV RNA (B) and neopterin (C) in patients with plasma HIV RNA < 20 copies/mL. “CSF”, cerebrospinal fluid; “CSF EBV+”, detectable CSF EBV DNA

## Discussion

In this study EBV DNA was detectable at low concentrations in up to 21% of HIV-positive patients and it was associated with higher CSF HIV viral load and up to three-time higher rate of pleocytosis. Such prevalence decreased in treated patients although in 13.4% of those with undetectable plasma viral load EBV was still detectable and associated with worse compartmental virological and immunological biomarkers.

Several studies reported the role of EBV in immunocompromised patients and they showed that the detection of EBV DNA in the CSF is a good marker of primary central nervous system lymphoma (PCNSL, with sensitivity and specificity of 70% and 80%) [19] [20] [21] [22]. Some studies reported the development of PCNSL in HIV-infected EBV-positive patients [23] [24] [25] [26].

Weinberg and coll. reported in some patients the presence of pleocytosis, detectable CSF EBV DNA and EBV related-mRNA supporting the hypothesis that EBV DNA is not carried by latently infected inflammatory cells but from actively replicating virus [27]. Furthermore EBV affects the immune system and it may enhance neuronal degeneration in chronic inflammatory conditions. [13][28]. Higher rates of B-amyloid protein and neurofibrillary tangles have been observed in the brains of EBV-positive patients diagnosed with Alzheimer’s dementia as compared to controls [29]. EBV may cause a sub-clinical chronic infection and facilitate inflammatory cells’ trafficking through the BBB thus increasing HIV entrance into the CNS. [30][31]. Our data seem to confirm this hypothesis since both naïve and treated patients present higher CSF HIV viral loads (and CSF to plasma HIV RNA ratios) and white blood cells. Additionally cART-treated patients with detectable CSF EBV DNA showed higher CSF to albumin ratios supporting a potential role in the persistence of BBB damage. The latter has been shown to be a prevalent feature of patients with dementia, to persist in some subjects despite treatment and to be associated with markers of neuronal damage and inflammation [32][33][34]. Hosting EBV astrocytes, and microglia, theoretically could create a wide net of cell-to-cell crosstalk encouraging migration of monocytic/macrophagic line cells and modifying CNS physiological homeostasis [35][36][37]. This effect may be independent from HIV control and immune system improvement: these conditions have been associated with the absence of neuronal damage and with the lowest CSF concentrations of neopterin [38][39][40].

In a recent study that analyzed 108 gut biopsies collected from 19 HIV-infected and 22 HIV-uninfected participants, CMV and EBV were detected in more than 70% of samples but more commonly in HIV-positive subjects [41]. While the negative effects of sporadic or continuous CMV replication are well-known, there is still uncertainty on the role of EBV in chronic immune activation.

Additionally EBV may have a role in suppressing the CNS immune system and therefore maintain an incomplete T-cell mediated inflammatory response; this may be achieved through the expression of viral genes encoding for proteins with immunoevasin-like function. This may translate into higher rates of pleocytosis but with less inflammatory activity [42][43]. On the other hand our EBV-positive treated patients showed lower CD4 counts thus suggesting that immune control may be needed in order to restore a partial control on EBV low level replication.

Some limitations of this study should be acknowledged including the low sample size, the lack of a control group and the lack of plasma EBV DNA measurements. Additionally our cohorts include several patients with very low nadir CD4 cell counts and heterogeneous clinical conditions: the same effect may not be observed in less advanced individuals.

In conclusion we reported for the first time the prevalence of EBV detection in the CSF of HIV-positive patients without lympho-proliferative disorders. Besides we observed that naïve subjects with detectable CSF EBV DNA had a higher HIV viral load and higher markers of neuronal damage and inflammation; in treated individuals despite a higher HIV viral load we report a higher prevalence of blood brain barrier damage, pleocytosis and immune activation. Further studies are warranted for understanding the contribution of EBV to HIV-associated CNS disorders.

